# ZEB1, a novel regulator of Junctional Adhesion Molecule A, impacts sensitivity of pancreatic cancer-associated fibroblasts to oncolytic reovirus

**DOI:** 10.1101/2025.03.14.643289

**Authors:** Nicole Dam, Tom J. Harryvan, Hao Dang, Gavriil Ioannidis, Bernhard Schmierer, Lukas J.A.C. Hawinkels, Vera Kemp

## Abstract

Oncolytic virus (OV) therapy is a promising treatment for various tumors. However, in pancreatic ductal adenocarcinoma (PDAC), the high abundance of cancer-associated fibroblasts (CAFs) can limit OV therapy efficacy by impairing viral spread and anti-tumor immunity. We have previously shown that oncolytic reovirus infection of CAFs depends on expression of the reovirus entry receptor Junctional Adhesion Molecule A (JAM-A), which is not or lowly expressed in most PDAC CAFs. We propose that increasing JAM-A expression on CAFs will boost viral spread in a tumor. However, there are currently no known regulators of JAM-A expression.

Therefore, we performed a genome-wide CRISPR/Cas9 knock-out screen to identify regulators of JAM-A expression. Ablation of the top negative regulator, Zinc Finger E-Box binding Homeobox 1 (*ZEB1*), in pancreatic fibroblasts led to strong JAM-A upregulation. We show that ZEB1 directly regulates JAM-A expression by binding to the E-box regions located within the JAM-A promotor. Importantly, ZEB1 ablation increased the sensitivity of fibroblasts to reovirus infection and subsequent cell death. Our work provides a novel overview of genes regulating JAM-A expression and provides a rational approach of combining ZEB1 inhibition with reovirus therapy to target both CAFs and tumor cells in stroma-rich tumors such as PDAC.

## Introduction

Oncolytic viruses (OVs) are a novel anti-cancer therapy currently tested in pre-clinical research and clinical trials for various tumor types, including pancreatic cancer.^(1, 2)^ The anti-tumor activity of OVs is attributed to a dual mechanism, relying on both direct killing of tumor cells and indirect activation of an anti-tumor immune response through release of immune-stimulating molecules in the tumor microenvironment (TME).^(3)^ OVs currently tested in (pre-)clinical studies have a tropism to cancer cells either intrinsically or upon genetic manipulation.^(4)^ However, for many tumor types, including pancreatic ductal adenocarcinoma (PDAC), the tumor-associated stroma constitutes up to 80% of the tumor mass, which could hamper the efficacy of oncolytic virotherapy.^(5, 6)^

An important and highly abundant cell type in the TME of pancreatic cancer is the cancer-associated fibroblast (CAF). CAFs are known to influence tumor development and progression, therapy sensitivity and anti-tumor immune responses.^(7)^ Among many other functions, CAFs are known to induce desmoplasia by producing extracellular matrix around the tumor. Desmoplasia has been thought to influence sensitivity to OVs by for example hindering OV penetration and spread in the tumor and by hampering immune cell infiltration.^(8, 9)^ Furthermore, CAFs can produce cytokines and chemokines that inhibit the activity of anti-tumor immune responses.^(10)^ Therefore, targeting CAFs with OVs, in addition to tumor cells, could be beneficial to increase overall therapy effectiveness.

While several papers have previously developed genetically modified OVs targeted to CAFs, ^(11-13)^ our group previously found that CAFs expressing the reovirus entry receptor Junctional Adhesion Molecule A (JAM-A) can be targeted by unmodified oncolytic reovirus.^(14)^ However, most CAFs in PDAC do not express JAM-A on their cell-surface and are therefore resistant to reovirus-induced cell death. Artificial introduction of JAM-A onto JAM-A negative fibroblasts sensitized fibroblasts to reovirus and increased reovirus infection rates in a co-culture model of tumor cells and fibroblasts.^(14)^ Strategies to increase JAM-A expression would thus increase the sensitivity of fibroblasts, and potentially the whole tumor, to reovirus induced cell death. However, it remains to be elucidated how JAM-A expression is regulated.

Therefore, in this study a CRISPR/Cas9 genome-wide knockout (KO) screen was performed to identify factors that regulate JAM-A expression. This screen identified Zinc Finger E-box binding Homeobox 1 (*ZEB1*) and Fibroblast Growth Factor Receptor 1 (*FGFR1*) as strongest negative regulators of JAM-A expression. Validation experiments showed that ablation of ZEB1 led to an induction of JAM-A expression, even on JAM-A negative cell lines. Mechanistically, ZEB1 was shown to bind directly to the JAM-A promotor causing its downregulation. Reovirus replication and apoptotic cell death were increased upon ZEB1 ablation in pancreatic fibroblasts, providing a rational approach to combine ZEB1 targeting with oncolytic reovirus to treat stroma-rich tumors like PDAC.

## Results

### A genome-wide CRISPR/Cas9 KO screen identifies novel regulators of cell surface JAM-A expression in pancreatic fibroblasts

Given the important role of JAM-A in mediating entry of reovirus into fibroblasts and its role in inducing virus-mediated apoptosis,^(14)^ we performed a genome-wide CRISPR/Cas9 KO screen to identify how JAM-A expression on the cell surface of fibroblasts is regulated. RLT-PSC, a pancreatic stellate cell line with moderate JAM-A expression levels, was used to identify both positive and negative regulators of JAM-A surface expression. Stable Cas9-expressing RLT-PSCs were transduced with a genome-wide sgRNA library ^(15)^ and the mutagenized cell population was subsequently sorted for JAM-A^high^ and JAM-A^low^ populations (**Figure 1A**). The integrated guide cassettes in sorted fibroblasts were then deep sequenced to determine the relative absence or enrichment of sgRNAs, resulting in an unbiased overview of genes involved in cell-surface expression of JAM-A (**Figure 1A-F**). Among the highly significant positive regulators, in which gene KO results in reduced JAM-A surface expression, F11 receptor (*F11R*) was the top enriched hit in the JAM-A^low^ population in both replicates of the screen. *F11R* is the gene encoding JAM-A, confirming the validity of the screen (**Figure 1B,C,E**). Another positive regulator found enriched in the JAM-A^low^ population was *SPPL3* (**Figure 1B,C,E)**, a gene that was recently described to control the composition of the cell surface glycosphingolipid (GSL) repertoire by inhibiting the glycosyltransferase B3GNT5.^(16)^

**Figure 1.**
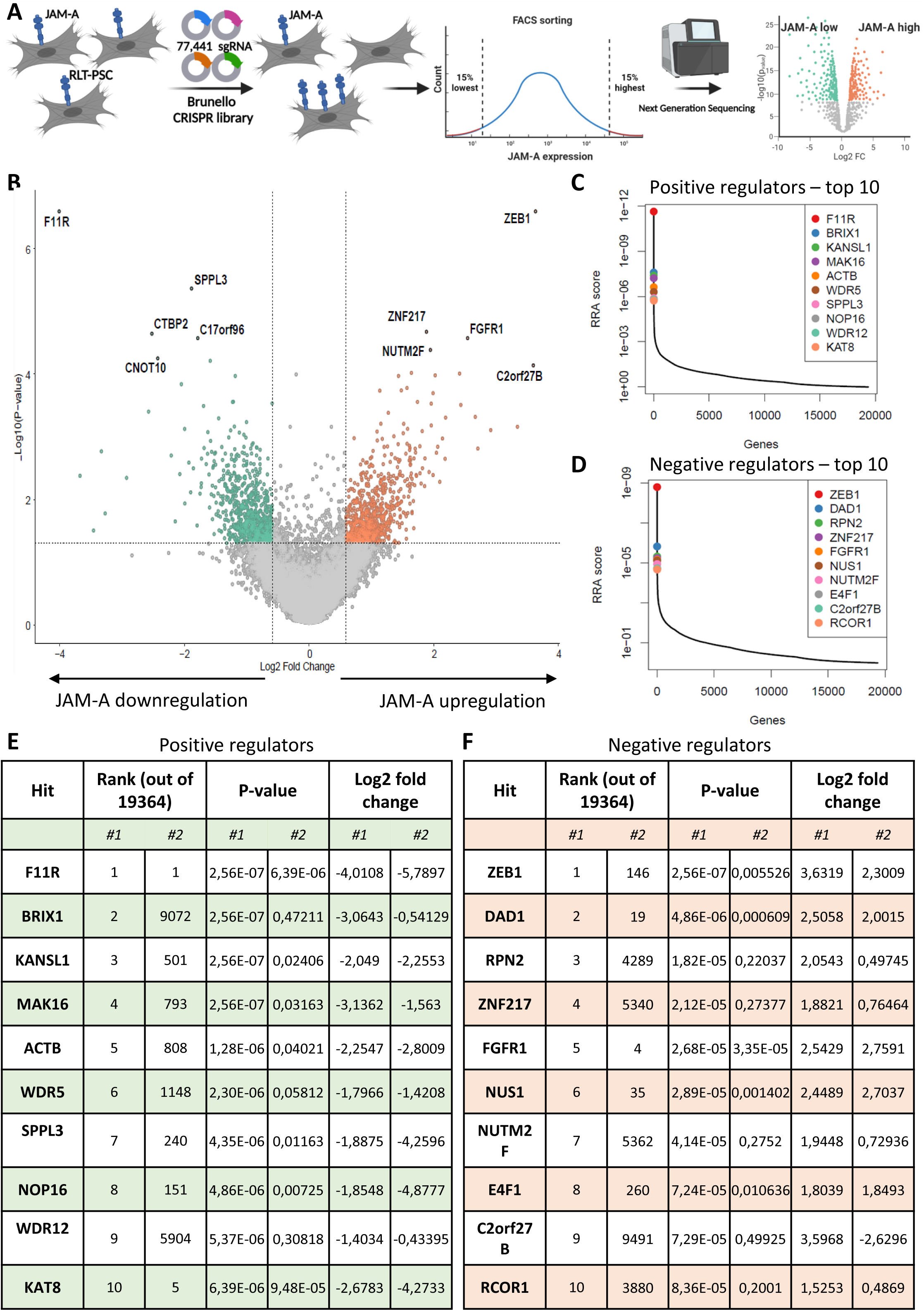
Genome wide CRISPR/Cas9 KO screening identifies positive and negative regulators of JAM-A expression on pancreatic fibroblasts. **A** Overview of the CRISPR/Cas9 screening approach to identify regulators of cell-surface JAM-A expression. Figure made with Biorender.com. **B** Volcano plot showing negative (green) and positive (orange) regulators of JAM-A. **C,D** The top 10 hits of positive (C) and negative (D) regulators of JAM-A cell surface expression. RRA score = robust rank aggregation score determined by MAGeCK computational tool. **E,F** Table displaying the top 10 hits displayed in C and D with corresponding rank, as determined by MAGeCK RRA score, P-values and log2 fold change, in both replicate 1 (left columns) and replicate 2 (right columns) of the screen.

The negative regulators of cell-surface JAM-A expression were deemed as targets of particular interest, because targeting these therapeutically could enforce stromal JAM-A expression and potentially boost reovirus activity in tumors. Two of the top hits in the JAM-A^high^ population in both replicates of the screen were Zinc Finger E-box binding Homeobox 1 (*ZEB1*, rank 1/19364 and rank 146/19364, respectively) and Fibroblast Growth Factor Receptor 1 (*FGFR1*, rank 5/19364 and rank 4/19364) (**Figure 1B,D,F**). ZEB1 is a transcription factor that is well described for its role in epithelial-to-mesenchymal transcription (EMT) and regulating expression of cell adhesion molecules like E-Cadherin.^(17)^ FGFR1 is part of the family of FGFRs, which together are responsible for a variety of biological processes, including cell growth, migration, differentiation, survival, and apoptosis.^(18)^

### SPPL3 KO results in shielding of JAM-A on the cell surface, but does not influence reovirus-mediated killing

Since SPPL3 KO was identified as a significant positive, and potentially targetable, regulator in the CRISPR/Cas9 screen, we further analyzed this hit. It has been shown that *SPPL3* KO in HAP1 cells leads to accumulation of negatively charged neolacto-series GSLs (nsGSLs) on the cell surface and this prevents HLA class I (HLA-I) interactions with immune receptors. This also leads to shielding of cell-surface HLA-I from being bound by an anti-HLA-I antibody.^(16)^ Since JAM-A is also a cell-surface molecule, we hypothesized that shielding also occurs for JAM-A. This could potentially affect the entry of reovirus into the cell and thereby susceptibility to reovirus-induced killing. To investigate this, we performed a flow cytometry-based antibody titration assay of JAM-A on HAP1 wildtype and HAP1 *SPPL3* KO cells. At lower antibody concentrations a clear decrease in JAM-A was observed following *SPPL3* KO, but JAM-A levels at higher antibody concentrations were almost similar between wildtype and *SPPL3* mutant cells, indicative of JAM-A shielding (**Figure 2A,B**). However, no apparent effect on subsequent susceptibility to reovirus-induced killing was observed between wildtype and *SPPL3* KO cells (**Figure 2C**) and therefore this target was not further pursued.

**Figure 2.**
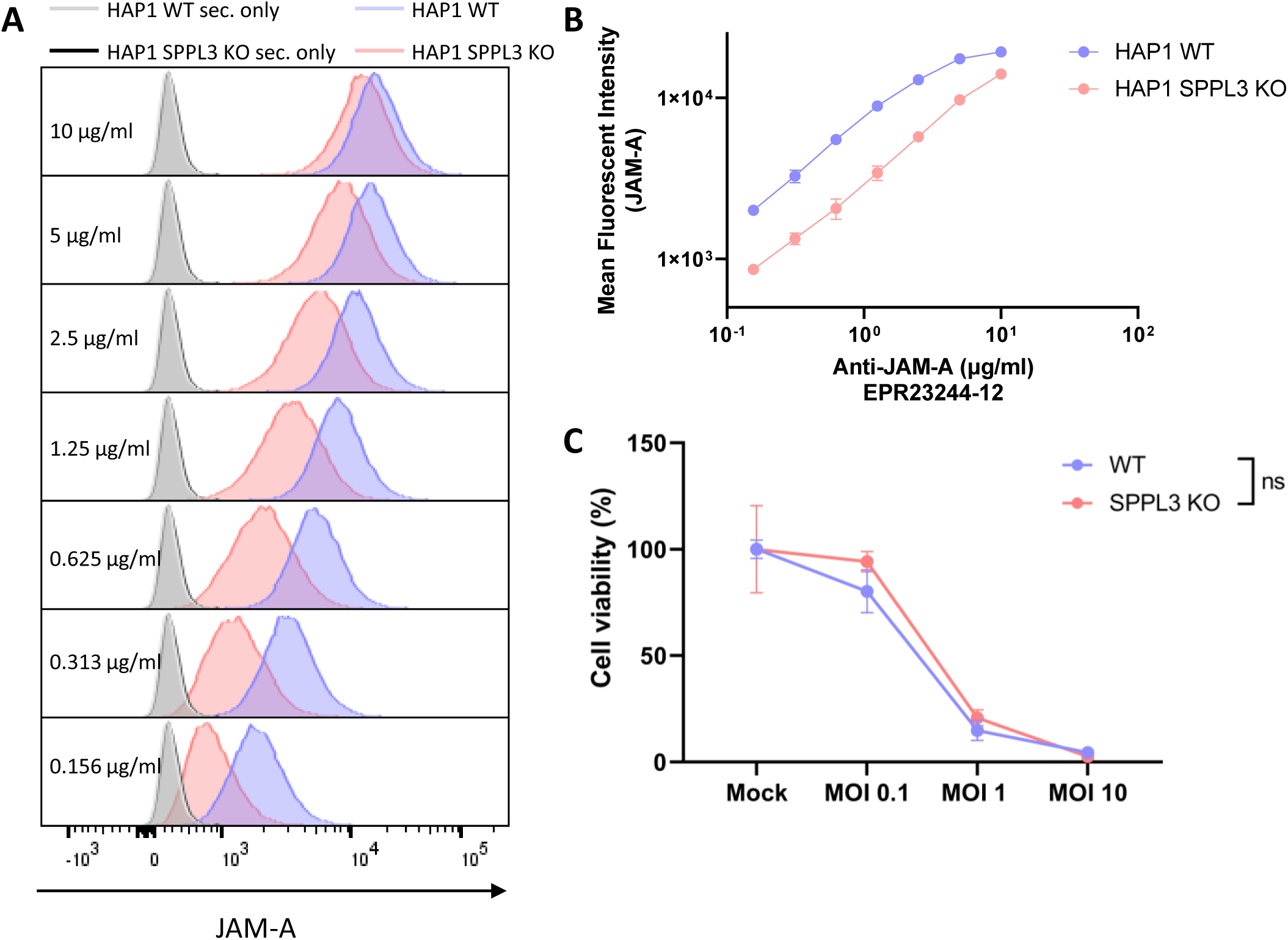
SPPL3 shields JAM-A from flow cytometric detection, but KO does not result in different reovirus sensitivity. **A** Flow cytometric titration of JAM-A antibody on HAP1 WT and HAP1 SPPL3 KO cells. Grey: secondary only antibody HAP1 WT, black: secondary only antibody HAP1 SPPL3 KO, blue: stained HAP1 WT, pink: stained HAP1 SPPL3 KO. **B** Mean fluorescent intensity of JAM-A at different concentrations of JAM-A antibody. Data is derived from a representative experiment and plotted as mean ± SD. **C** Cell viability (%) compared to mock of HAP1 WT and SPPL3 KO cells following three days reovirus infection at different MOIs, as determined by WST-1 assay. Significance was calculated using two-way ANOVA with correction for multiple testing (Šídák’s test), significance is depicted at MOI 10, ns = not significant. Data is derived from a representative experiment and plotted as mean ± SD.

### Validation of CRISPR/Cas9 screen derived negative regulators shows that knockdown of ZEB1 results in JAM-A upregulation

To verify whether the negative regulators found in the CRISPR/Cas9 screen are valid, ZEB1 and FGFR1 shRNA-mediated knockdowns (KDs) were generated in the RLT-PSC stellate cell line in which the CRISPR/Cas9 screen was performed and a murine pancreatic CAF that expresses low levels of JAM-A (KPC3-CAF1). Following verification of the KD (**Figure 3A-F**), we performed flow cytometric analyses of cell-surface JAM-A expression. FGFR1 KD did not result in upregulation of JAM-A, contrasting the results of the genome-wide CRISPR/Cas9 KO screen. However, partial shRNA-mediated ablation of ZEB1 resulted in a strong increase in JAM-A expression, both in JAM-A high (RLT-PSC) and low (KPC3-CAF1) expressing fibroblasts (**Figure 3G,H**), which is in line with the results of the CRISPR/Cas9 screen. Furthermore, we show that ZEB1 KD in both fibroblasts results in sensitization to reovirus-mediated cell death (**Figure 3I,J**). JAM-A upregulation and sensitization to reovirus upon ZEB1 KD, but not FGFR1, was also observed in the transformed skin fibroblast cell line NBS **(Figure S1A-E)**. All in all, this shows that ZEB1 is a potent negative regulator of cell-surface JAM-A expression, and KD of ZEB1 results in JAM-A upregulation in pancreatic, but also other, fibroblasts.

**Figure 3.**
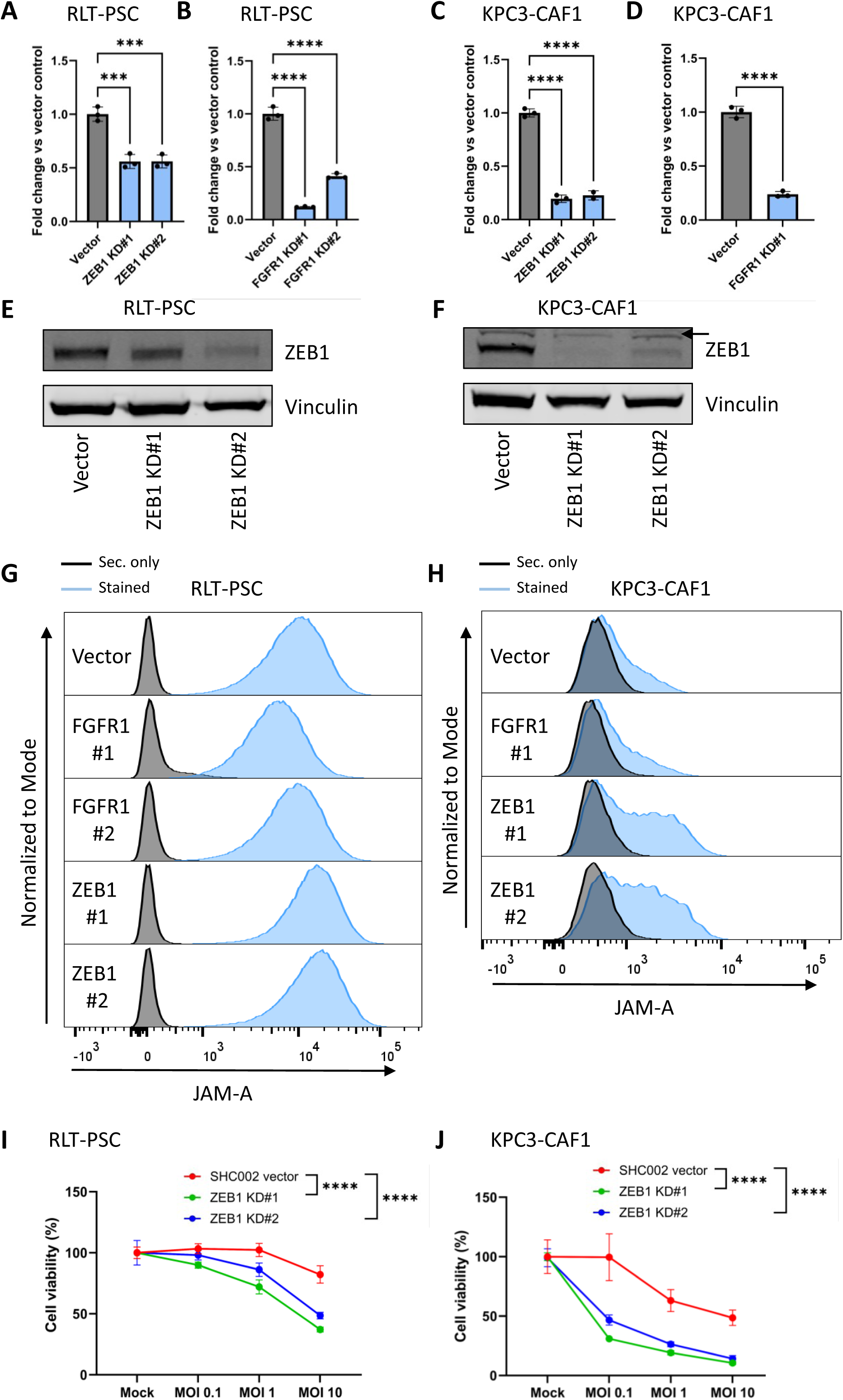
ZEB1 KD, but not FGFR1 KD, results in increased JAM-A expression in JAM-A positive and negative human and murine fibroblast cell lines and sensitization to reovirus. **A-D** RT-qPCR for *ZEB1* expression in RLT-PSC (A) and KPC3-CAF1 (C) vector control and ZEB1 KD and *FGFR1* expression in RLT-PSC (B) and KPC3-CAF1 (D) vector control and FGFR1 KD. Ct values were corrected for *IPO8* and *EIF2B1* (RLT-PSC) or *Mzt2* and *Ptp4a* (KPC3-CAF1) expression and calculated as fold change vs vector control. Significance was calculated using one-way ANOVA with correction for multiple testing (Šídák’s test) or unpaired T-test for KPC3-CAF1 FGFR1 KD, *** p≤0,001, ****p≤0,0001. Data is derived from a representative experiment and plotted as mean ± SD. **E,F** Western blot for ZEB1 with vinculin as loading control in RLT-PSC (E) and KPC3-CAF1 (F) vector control and ZEB1 KD. Arrow indicates aspecific band. **G,H** Flow cytometric analysis of cell-surface JAM-A expression in RLT-PSC (G) and KPC3-CAF1 (H) vector control, FGFR1 KD and ZEB1 KD. Black: secondary antibody only, blue: stained. **I,J** Cell viability (%) relative to mock following infection of RLT-PSC (J) and KPC3-CAF1 (K) vector control and FGFR1 and ZEB1 KDs with reovirus at multiple MOIs for 3 days, as measured by a WST-1 assay. Significance was calculated using two-way ANOVA with correction for multiple testing (Šídák’s test), significance is depicted at MOI 10, ****p≤0,0001. Data is derived from a representative experiment and plotted as mean ± SD.

### Complete ZEB1 ablation in JAM-A positive and negative human and murine fibroblast lines results in upregulation of cell-surface JAM-A expression through direct transcriptional regulation

Because of the strong effects of ZEB1 KD on JAM-A expression, we decided to further focus on ZEB1 as a potent negative regulator of JAM-A expression. To this end, we used CRISPR/Cas9 mediated targeting of a conserved region of the *ZEB1* exon 1-intron boundary in pancreatic fibroblasts of human (RLT-PSC, hPS1) and murine (KPC3-CAF1) origin. Clonal knockout (KO) cell lines were generated by single cell sorting and subsequently rescued by re-introducing *ZEB1* cDNA into the cells. *ZEB1* KO was validated on DNA level by PCR and Sanger sequencing of the region around the gRNA. Since sequencing was difficult due to high GC content of this region, PCR products were transformed in bacteria, isolated and sequenced. This showed that indels that cause a frameshift or are large enough to disrupt the protein had occurred in all KO clones (**Figure S2A-D**). Interestingly, we could observe many different edits in the RLT-PSC KO cell lines (**Figure S2B**), which can be explained by the fact that RLT-PSC has a mean number of 60 chromosomes per cell ^(19)^ and thus potentially multiple *ZEB1* alleles per cell. For hPS1, 2 different indels per KO cell line could be identified, providing proof of clonality of these lines (**Figure S2D**). Furthermore, KO and rescue of ZEB1 protein was validated using western blot (**Figure 4A-C**).

**Figure 4.**
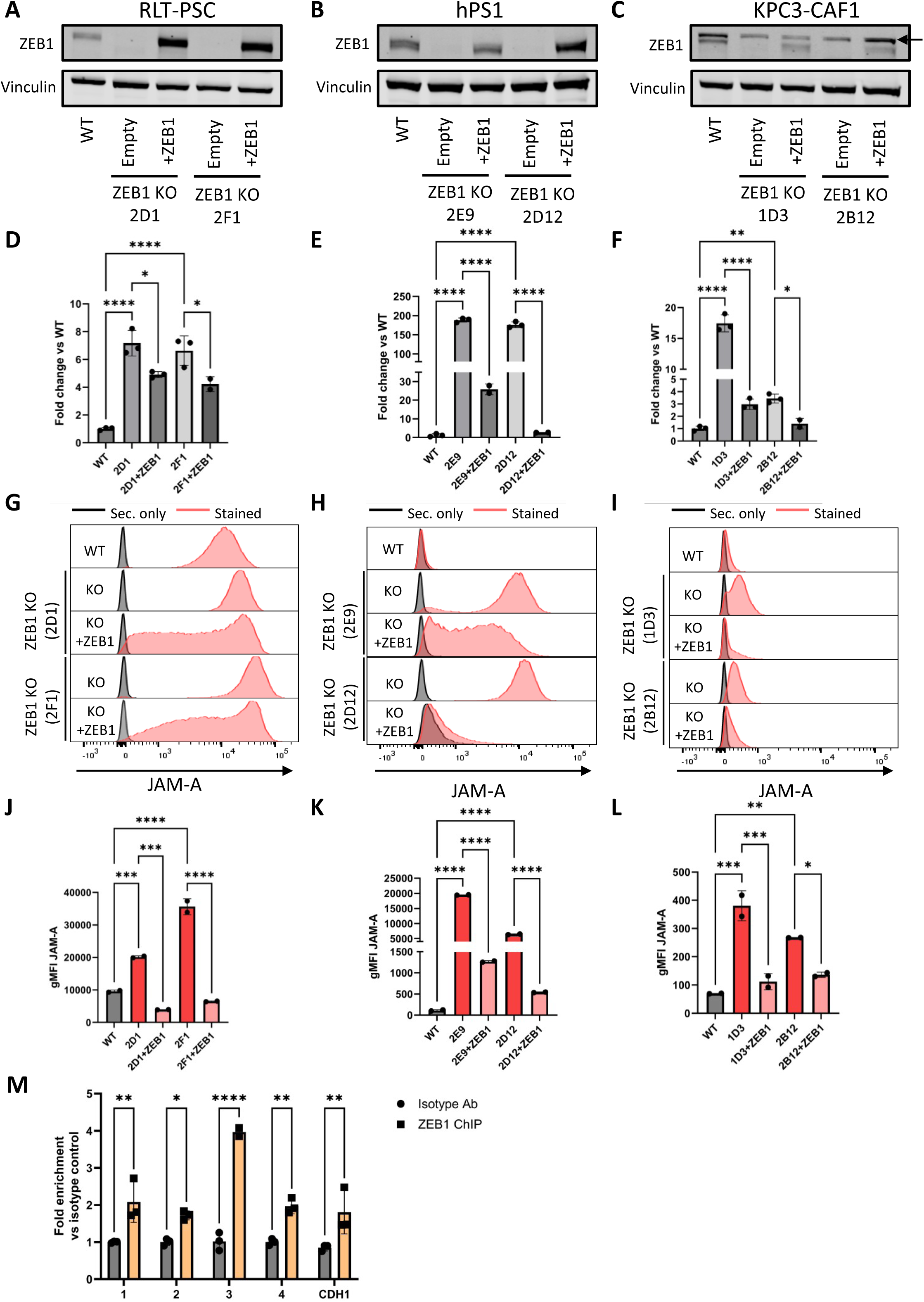
ZEB1 KO in human and murine pancreatic fibroblasts results in increased JAM-A RNA and protein through direct transcriptional regulation. **A,B,C** Western blot of clonal ZEB1 KO and ZEB1 rescues in RLT-PSC (A), hPS1 (B) and KPC3-CAF1 (C) fibroblasts with vinculin (vinc) as loading control. Arrow indicates aspecific band. **D,E,F** qRT-PCR analysis of JAM-A expression in ZEB1 KO and rescue cell lines RLT-PSC (D), hPS1 (E) and KPC3-CAF1 (F). Ct values were corrected for *IPO8* and *EIF2B1* (RLT-PSC and hPS1) or *Mzt2* and *Ptp4a2* (KPC3-CAF1) expression and calculated as fold change vs WT. *p≤0,05, **p≤0,01, ****p≤0,0001 as determined by one-way ANOVA with correction for multiple testing (Šídák’s test) Data is derived from a representative experiment and plotted as mean ± SD. **G,H,I** Flow cytometric analyses of cell-surface JAM-A expression of ZEB1 KO clones and rescues in RLT-PSC (G), hPS1 (H) and KPC3-CAF1 (I). Black: secondary only antibody, red: stained. **J,K,L** Geometric mean fluorescent intensity (gMFI) of JAM-A expression of the different cell lines as depicted in G,H,I. *p≤0,05, **p≤0,01, ***p≤0,001, ****p≤0,0001 as determined by one-way ANOVA with correction for multiple testing (Šídák’s test). Data is derived from a representative experiment and plotted as mean ± SD. **M** Chromatin immunoprecipitation qPCR (ChIP-qPCR) assay to identify DNA binding regions of ZEB1. 1-4: different E-box binding regions within the F11R promotor, CDH1: E-box binding region within the E-cadherin promotor. *p≤0,05, **p≤0,01, ****p≤0,0001 as determined by two-way ANOVA with correction for multiple testing (Šídák’s test). Data is derived from a representative experiment and plotted as mean ± SD.

qRT-PCR analysis revealed that ZEB1 ablation resulted in significant upregulation of JAM-A RNA levels (**Figure 4D-F**). The extent of JAM-A upregulation varies within each clone, which can be attributed to the polyclonal nature of the WT cell lines.^(19-21)^ Reintroducing *ZEB1* cDNA caused a significant downregulation of JAM-A RNA expression for all lines compared to their corresponding KO clone (**Figure 4D-F**). The changes in RNA expression were also validated on protein level through flow cytometric analyses of cell-surface JAM-A expression. RLT-PSC pancreatic fibroblasts already express a moderate level of JAM-A, but still showed a strong upregulation of JAM-A following ZEB1 KO (**Figure 4G,J**). The most striking phenotype was observed in hPS1 (**Figure 4H,K**) and the mouse pancreatic CAF KPC3-CAF1 (**Figure 4I,L**), in which ZEB1 KO transforms the cells from JAM-A negative to strongly positive. For all cell lines, reintroduction of *ZEB1* cDNA into the KOs results in a significant downregulation of JAM-A cell-surface expression (**Figure 4H-L**). Since ZEB1 is known to regulate the expression of integrin β1,^(22)^ a known secondary entry receptor for reovirus,^(23)^ we checked whether there are changes in integrin β1 following ZEB1 KO that could influence potential differences in reovirus infection. However, no differences were found in integrin β1 expression in the ZEB1 KO clones of both RLT-PSC and hPS1 fibroblasts compared to WT (**Figure S3**).

Since ZEB1 is known to bind directly to E-box regulatory regions within the E-cadherin (CDH1) promotor region,^(24, 25)^ we hypothesized that it also binds to E-box regions within the *F11R* (JAM-A) promotor to regulate its expression. Using a chromatin immunoprecipitation qPCR (ChIP-qPCR) assay with primers targeting different E-Box sequences within the *F11R* promotor (numbered 1 to 4), we validated that ZEB1 is more enriched in binding to the *F11R* promotor when compared to isotype control antibody-bound beads. ZEB1 bound to a similar extent to the *F11R* promotor as to the CDH1 promotor (**Figure 4M**). This demonstrates that ZEB1 can inhibit JAM-A expression by directly binding to the *F11R* promotor, explaining the increased levels of both JAM-A RNA and protein following ZEB1 KO.

Altogether, these data show that ZEB1 is a novel, strong regulator of JAM-A in human and murine pancreatic fibroblasts and CAFs and that ablation of this transcription factor enhances the expression of JAM-A through direct transcriptional regulation, even in fibroblasts that do not express JAM-A under basal conditions.

### ZEB1 KO in pancreatic fibroblasts and CAFs increases reoviral replication and sensitizes to reovirus-induced apoptotic cell death

Having established that ZEB1 is a potent negative regulator of cell-surface JAM-A expression on pancreatic fibroblasts and CAFs, we next aimed to confirm whether ZEB1 ablation would result in increased susceptibility to reovirus infection and reovirus-mediated cell death. ZEB1 KO pancreatic fibroblasts were exposed to different reovirus concentrations, followed by assessment of cell viability, caspase 3/7 activation to determine activation of the apoptotic pathway and assessment of reovirus protein σ3 levels in the cell as a measure for viral infection and replication. While RLT-PSC WT, due to its moderate JAM-A expression, can be killed by reovirus, ZEB1 KO significantly increased reovirus-induced cell death at high MOI (**Figure 5A**). Furthermore, we show that this increased level of cell death is mediated through apoptotic cell death (**Figure 5B,C, supplemental video**). In addition to increased levels of cell death, we also observed increased levels of reovirus σ3 protein in ZEB1 KO clone 2D1, indicative of increased reovirus replication (**Figure 5D**). Of note, while clone 2F1 did not show increased σ3 levels (**Figure 5D**), JAM-A expression (**Figure 4G**) and cell death (**Figure 5A**) were strongly increased. While hPS1 WT is not susceptible to reovirus, ZEB1 KO in this cell line strongly increased both reovirus replication and apoptotic cell death (**Figure 5E-H, supplemental video**). Similarly, ZEB1 KO in the murine CAF line KPC3-CAF1 resulted in higher levels of viral replication, reovirus-induced cell death and caspase 3/7 activation (**Figure 5I-L, supplemental video**). Finally, restoring ZEB1 expression in the ZEB1 KO clones either completely or partially restored the resistance to reovirus-mediated cell death (**Figure 5A,E,I**).

**Figure 5.**
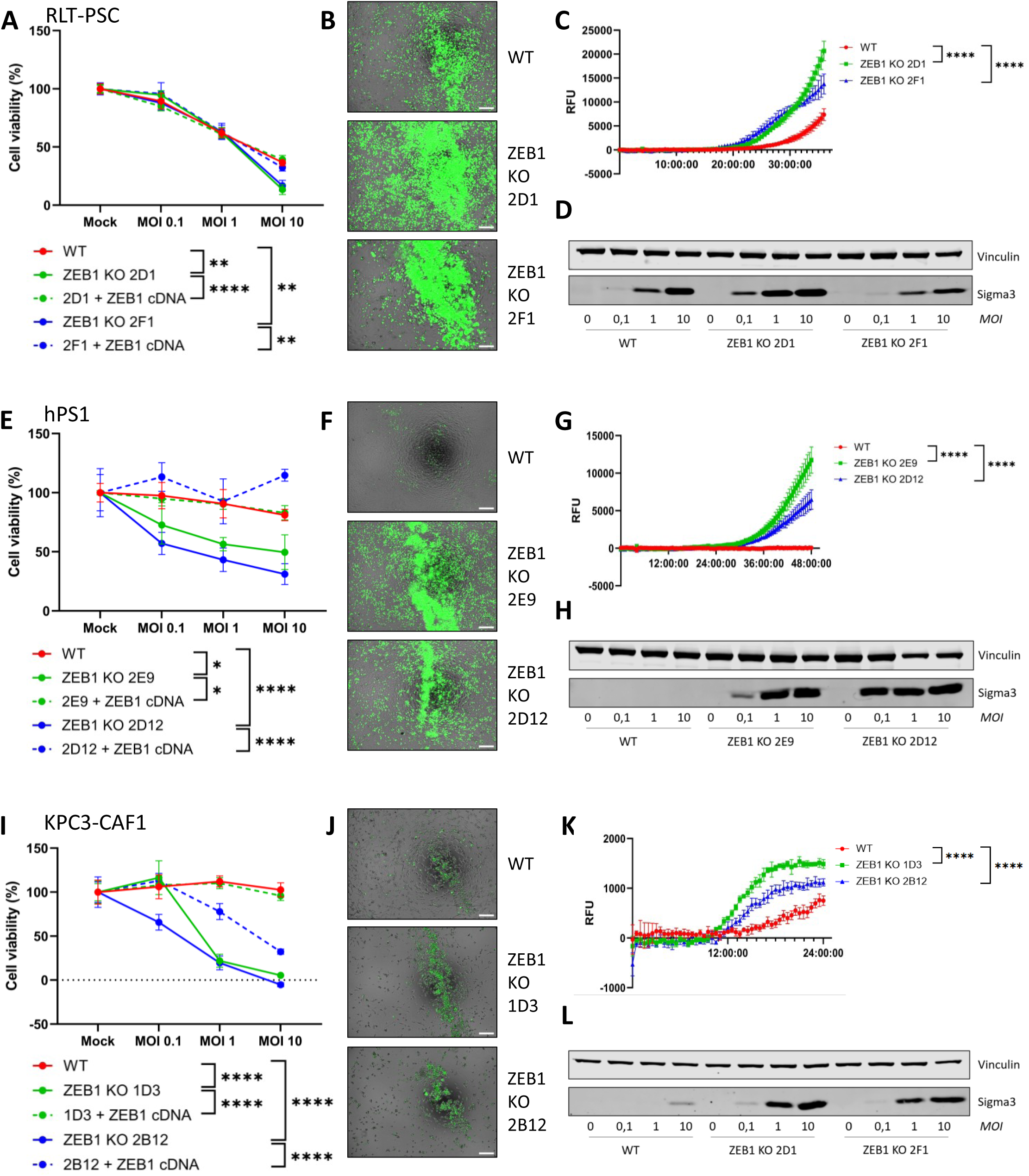
ZEB1 KO in human and murine fibroblasts results in increased susceptibility to reovirus infection and reovirus-mediated apoptotic cell death. **A,E,I** Cell viability (%) relative to mock following infection with reovirus at multiple MOIs, as measured by a WST-1 assay. RLT-PSC (A) were infected for 4 days, hPS1 (E) for 6 days and KPC3-CAF1 (I) for 3 days. Significance was calculated using two-way ANOVA with correction for multiple testing (Šídák’s test), significance is depicted at MOI 10, *p≤0,05, **p≤0,01, ****p≤0,0001. Data is derived from a representative experiment and plotted as mean ± SD. **B,F,J** Overlay of phase-contrast and GFP images of CellEvent Caspase3/7 assay of RLT-PSC (B), hPS1 (F) and KPC3-CAF1 (J) WT and ZEB1 KO cells. Cells were infected with R124 MOI 10 for 36, 48 and 24 hours, respectively. Scale bar: 200 µM. **C,G,K** Quantification of CellEvent Caspase3/7 fluorescent signal over time, during infection with reovirus R124 MOI 10 for 36 (RLT-PSC, C), 48 (hPS1, G) or 24 hours (KPC3-CAF1, K). Significance was calculated using two-way ANOVA with correction for multiple testing (Šídák’s test), ****p≤0,0001. Data is derived from a representative experiment and plotted as mean ± SD. **D,H,L** Western blot for reovirus Sigma3 expression and vinculin as loading control in WT and ZEB1 KO cells following reovirus infection at different MOIs. RLT-PSC (B) and hPS1 (E) were infected for 2 days and KPC3-CAF1 (H) for 1 day.

All in all, this shows that ZEB1 ablation in pancreatic fibroblasts, in addition to causing JAM-A upregulation, can increase their susceptibility to reovirus infection and reovirus-mediated apoptotic cell death.

## Discussion

In this study, we identified ZEB1 as a key factor regulating the expression of the reoviral entry receptor JAM-A, using a genome-wide CRISPR/Cas9 KO screen in fibroblasts. Clonal ZEB1 KOs and KDs in three different pancreatic fibroblast cell lines indeed showed an upregulation of JAM-A cell-surface expression and subsequent sensitization to reovirus replication and reovirus-mediated cell death. Mechanistically, we show that ZEB1 binds directly to E-box regulatory regions located within the JAM-A promotor, thereby repressing its transcription.

The field of oncolytic virotherapy has, until recently, focused on finding and developing tumor-selective viruses that leave healthy cells intact.^(26)^ While this improves safety, it can also limit efficacy by preventing targeting of other components in the TME. In several tumor types, including PDAC, a large part of the tumor stroma consists of CAFs,^(5)^ which can form a barrier to the efficacy of OV therapy. Therefore, strategies to also target the PDAC stroma, such as presented here, hold great potential to improve the therapeutic efficacy of the viruses.

Several groups have focused on generating genetically modified OVs directed against CAFs.^(11-13)^ Intriguingly, we previously found that reovirus inherently has the capacity to target stromal components as long as they express the reovirus entry receptor JAM-A.^(14)^ Reovirus is particularly interesting for extending its tropism because it is non-pathogenic and causes a self-limiting infection in humans,^(27)^ in contrast to other OVs.^(28)^ We observed that reovirus-sensitive CAFs boosted overall viral spread in a multicellular tumor model compared to reovirus-resistant CAFs.^(14)^ Importantly, the majority of pancreatic CAFs express very low levels of JAM-A. Two approaches to boost the CAF tropism of reovirus could be: 1) modulating the viral vector to allow JAM-A independent entry ^(29)^ and 2) modulating the stroma to allow wildtype reoviral infection. Since JAM-A is crucial for induction of reovirus-mediated apoptosis,^(14)^ JAM-A independent entry could potentially interfere with the cytolytic capacity of these viruses. Furthermore, we found that there is also a strong correlation between the JAM-A expression on fibroblasts and cell death induced by our JAM-A independent reovirus Jin-3, showing that in the presence of JAM-A Jin-3 will still use this receptor to enter cells more efficiently.^(14)^ Therefore, modulating the stroma to increase reovirus infection of CAFs in a JAM-A dependent manner seems an attractive alternative. In order to increase the overall efficacy of reovirus therapy in pancreatic tumors, we propose to increase CAF targeting through upregulating JAM-A on their cell-surface.

Although the initial screen also identified *FGFR1* as a potent negative regulator of JAM-A expression, FGFR1 KD in both the RLT-PSC fibroblast cell line in which the screen was performed, and the murine CAF line KPC3-CAF1, did not result in JAM-A upregulation. Since the other hit, *ZEB1*, did show a strong JAM-A upregulation, this factor was further pursued. ZEB1 has been described as one of the main transcription factors regulating epithelial-to-mesenchymal transition (EMT). Furthermore, it regulates the expression of various cell-adhesion molecules, including E-Cadherin.^(17)^ Since JAM-A is also a cell-adhesion molecule like E-Cadherin, it is interesting to find that expression of both proteins is regulated by ZEB1.^(18)^ ZEB1 suppresses E-Cadherin expression by binding to enhancer-box (E-box) regulatory motifs,^(24, 25)^ and we show here that ZEB1 also binds to several E-box regulatory motifs present in the promotor of the *F11R* gene, which encodes JAM-A. Although ZEB1 is highly expressed in CAFs, tumor cells that undergo EMT, a common phenomenon during tumor progression, also upregulate ZEB1.^(30, 31)^ Loss of JAM-A in tumor cells during tumor progression has been observed in PDAC patients and is associated with adverse clinical outcome.^(32)^ Therefore, targeting of ZEB1 could potentially also restore JAM-A expression in these mesenchymal tumor cells, sensitize them to reovirus infection and perhaps even revert EMT.

Interestingly, a recent paper also showed important roles of ZEB1 expression in CAF polarization, promoting myofibroblastic features and restricting immune activation. ZEB1 ablation in fibroblasts of mice bearing colorectal cancer impaired the barrier function of CAFs by decreasing collagen deposition. Furthermore, it increased cytokine production by CAFs, which led to lymphocyte attraction and effective anti-tumor immune responses.^(33)^ Similar effects were observed in a murine model of breast cancer.^(34)^ Both effects of ZEB1 ablation in fibroblasts, decreased barrier function and increased immune activation, point to a strong synergistic anti-tumor effect when combined with reovirus therapy, increasing both viral spread and enhancing reovirus-induced anti-tumor immunity. Additionally, ZEB1 has also been shown to have anti-tumor effects in other components of the TME by mediating IL-2 suppression in tumor-specific T-cells. Therefore, inhibiting ZEB1 activity will restore IL-2 expression in T cells, resulting in increased T cell proliferation and homeostasis.^(35)^Thus, in addition to sensitizing CAFs to reovirus, there are multiple rationales why targeting ZEB1 in the PDAC TME, together with reovirus treatment, is an attractive treatment option.

In conclusion, using unbiased CRISPR/Cas9 genome-wide KO screening, we identified previously unknown regulators of cell-surface JAM-A expression on CAFs, with ZEB1 as potent negative regulator. This will allow the design of rational combination treatments targeting ZEB1 to convert naturally resistant PDAC CAFs to a reovirus-susceptible stroma, which will allow concurrent targeting of both tumor cells and CAFs.

## Materials and Methods

### Primary fibroblast, cell line and organoid culture

KPC3-CAF1 ^(21)^ and the pancreatic stellate lines hPS1 (kindly provided by H. Kocher, University of London, London, England) ^(20)^ and RLT-PSC ^(19)^ were maintained in Dulbecco’s modified Eagle’s medium (DMEM)/F12 (Thermo Fisher Scientific, Leiden, the Netherlands), supplemented with 8% fetal calf serum (FCS), 100 IU/ml penicillin and 100 μg/ml streptomycin (all Thermo Fisher Scientific). HAP1 wildtype and SPPL3 KO cells (kind gift of dr. M. Jongsma, Dept. Cell & Chemical Biology, LUMC, the Netherlands) ^(16)^ were cultured in DMEM supplemented with 8% FCS, 100 IU/mL penicillin and 100 μg/mL streptomycin (all Thermo Fisher Scientific). All cells were cultured at 37°C and 5% CO_2_ and routinely confirmed to be negative for mycoplasma contamination.

### Oncolytic viruses

The wild-type type 3 Dearing (T3D) reovirus strain R124 was isolated by plaque purification from a heterogenous T3D stock obtained from ATCC (VR-824), and propagated and purified as described previously.^(29)^ Infections of cells with R124 were performed in the standard culture medium with 2% FCS.

### Flow cytometry and fluorescence-assorted cell sorting (FACS)

For cell surface staining, cells were harvested and washed twice with FACS buffer, consisting of PBS/0.5% bovine serum albumin (BSA, Sigma-Aldrich, Amsterdam, the Netherlands) and 0.05% sodium azide (Pharmacy LUMC, Leiden, the Netherlands). Human fibroblasts were incubated with rabbit anti-human JAM-A (EPR23244-12, Abcam, Cambridge, UK) at 4°C for 45 minutes. Subsequently, cells were washed twice with FACS buffer and incubated with goat anti-rabbit-PE (Jackson ImmunoResearch Europe Ltd, United Kingdom) at 4°C for 45 minutes. For murine samples, cells were incubated with a directly AF488-conjugated rat anti-mouse JAM-A (H202-106, Bio-Rad laboratories, Nazareth, Belgium) at 4°C for 45 minutes. For integrin beta-1 (CD29) staining, human fibroblasts were incubated with a PE-conjugated mouse anti-human CD29 (MAR4, BD biosciences, Drachten, the Netherlands) at 4°C for 45 minutes. Samples were measured on a LSR-II flow cytometer (BD biosciences) and data was analyzed with FlowJo software, v10.6.1 (BD biosciences). For single-cell sorting of ZEB1 KO clones, an Aria FACS sorter (BD Biosciences) was used. For cell sorting of RLT-PSC fibroblasts in the CRISPR/Cas9 KO screen, cells were sorted using a Sony SH800 cell sorter (Sony Biotechnology Inc, San Jose, CA, USA).

### Genome-wide CRISPR/Cas9 Knockout library screen

#### Generation of stable Cas9 expressing fibroblasts

RLT-PSC fibroblasts were lentivirally transduced with pLenti-Cas9-T2A-Blast-BFP (Addgene #196714) to express a codon optimized, WT SpCas9 flanked by two nuclear localization signals linked to a blasticidin-S-deaminase – mTagBFP fusion protein via a self-cleaving peptide. Following blasticidin selection, a stable BFP+ population was isolated by repeatedly sorting for BFP expression until >90% Cas9+/BFP+ was obtained.

#### Guide library

The genome-wide Brunello single guide RNA (sgRNA) library ^(15)^ was synthesized as 79 bp long oligos (CustomArray, Genscript). The oligo pool was double-stranded by PCR using ds_Ultramer and amplification with primers ds_fw and ds_rev (**Table S1**) to include an A-U flip in the tracrRNA,^(36)^ 10 nucleotide long random sequence labels (RSLs), and an i7 sequencing primer binding site.^(37)^ The resulting PCR product (**Table S1**) was cloned by Gibson assembly into pLenti-Puro-AU-flip-3xBsmBI (Addgene #196709).^(37)^ The plasmid library was input sequenced to confirm representation and packaged into lentivirus in HEK-293T (ATCC) using plasmids psPAX2 (a gift from Didier Trono, Addgene #12260) and pCMV-VSV-G (a gift from Bob Weinberg, Addgene #8454). The virus-containing supernatant was concentrated with Lenti-X concentrator (Takara, Saint-Germain-en-Laye, France), aliquoted and stored in liquid nitrogen.

#### Library virus titration and large-scale transduction

The functional titer of the library virus was estimated from the fraction of surviving RLT-PSC cells after transduction of target cells with different amounts of virus and puromycin selection. For the screen, Cas9-BFP-expressing target cells were transduced with the library virus in duplicate at an approximate MOI of 0.3 and a coverage of 500-1,000x (500-1,000 cells per guide) in the presence of 2 µg/ml polybrene. Transduced cells were selected with 2 µg/ml puromycin from day 2 to day 6 post transduction. A control sample worth 80 million cells per replicate was harvested at day 5 post transduction. Cell numbers per replicate were kept at >= 80 million/replicate throughout to ensure full library coverage.

At day 7 post transduction, RLT-PSC were stained as described above and two-way FACS-sorted based on the expression of JAM-A, with collection of the lowest (15%) and highest (15%) JAM-A expressing cells. Collected cell pellets were stored at -20°C until extraction of genomic DNA.

#### Genomic DNA, Library preparation and NGS sequencing

Genomic DNA was isolated using the QIAamp DNA Blood Maxi or Mini Kit (Qiagen, Venlo, the Netherlands), and guide and UMI sequences were amplified in a three-step PCR protocol as described,^(37)^ using the primers given in **Table S1**. The amplicon was sequenced on Illumina NovaSeq6000, reading 20 cycles Read 1 with custom primer CRISPRSeq (**Table S1**); 10 cycles index read i7 to read the UMI, and six cycles index read i5 for the sample barcode. NGS data was analyzed with the MaGeCK software, v.0.5.6.^(38)^

### Lentiviral transductions and transgenic cell lines

For all lentiviral constructs, third-generation packaging vectors and HEK293T cells were used for the generation of lentiviral particles.^(39)^ To generate ZEB1 KO fibroblast cell lines (hPS1, RLT-PSC & KPC3-CAF1), a sgRNA, 5’-caccgCACTCACCGTTATTGCGCCG-3’ (lowercase nucleotides are compatible with the restriction site) targeting a conserved region of the *ZEB1* exon 1-intron boundary was cloned into *BsmBI*-digested plentiCRISPRv2-hygromycin (Addgene: #98291). This design enables targeting of both murine and human *ZEB1* with the same construct. Lentiviral particles were generated and after transduction, hygromycin-resistant fibroblasts were sorted, based on their gain of JAM-A expression (highest 5%), at single cell density and subsequently expanded to acquire clonal lines. Cells were selected with 200 µg/ml of hygromycin B (Merck). Knockout verification of clones was performed via western blot and DNA sequencing.

Rescue of the ZEB1 KO was performed by reintroducing human *ZEB1*-encoding cDNA, which was amplified from a fibroblast cDNA library using primers 5’-gatcctcgagaccATGGCGGATGGCCCC-3’ (forward, *XhoI* restriction site) and 5’-gatcaccggtTTAGGCTTCATTTGTCTTTTC-3’ (reverse, *AgeI* restriction site) using the Phusion High-Fidelity PCR Kit according to manufacturer’s instructions (Thermo Fisher Scientific). The resulting PCR product was gel purified using the NucleoSpin Gel and PCR Clean-up kit (Macherey-Nagel, Dueren, Germany) and 3′ A-overhangs were added by incubation with *Taq-*polymerase (DreamTaq Green PCR Master Mix (Thermo Fisher Scientific)) for 30 minutes at 72°C. This product was cloned into the pCR™4-TOPO® TA vector using the TOPO® TA Cloning® kit according to manufacturer’s instructions (Thermo Fisher Scientific). Subsequently, this clone was fully sequenced and confirmed to be *ZEB1* transcript variant 2 (CCDS7169). Lastly, this cDNA was subcloned into pLV-CMV-puromycin ^(14)^ and termed pLV-CMV-*ZEB1*-puromycin. For hPS1, since these already contain a puromycin resistance cassette, *ZEB1* cDNA was cloned into pLV-CMV-neomycin. Clonal ZEB1 KO cells were transduced with this vector and selected and subsequently cultured with puromycin (2 µg/ml; Sigma-Aldrich) or G418 (400 µg/ml; Thermo Fisher Scientific). Finally, ZEB1 rescue was confirmed through Western blot.

Knockdown constructs were acquired from the Mission TRC shRNA library (Sigma-Aldrich), with target sequences that are shown in **Table S2**. Cells were selected and cultured with 2 µg/ml puromycin (Sigma-Aldrich).

### RNA isolation and qRT-PCR analysis

Total RNA was isolated using the NucleoSpin RNA isolation kit (Macherey-Nagel, Düren, Germany) according to manufacturer’s instructions. cDNA was synthesized using SuperScript™ II Reverse Transcriptase (Invitrogen, Thermo Fisher Scientific), followed by RT-qPCR analysis using SYBR Green Master mix (Bio-Rad) and the iQ5 Multicolour Real-Time PCR Detection System (Bio-Rad). Target genes were amplified using specific primers (**Table S3**). The ΔΔCt method was applied to calculate the levels of gene expression, relative to a control condition.

### DNA sequencing of ZEB1 KO clones

To verify knockout on a genomic level, DNA was isolated using the Purelink gDNA mini kit (Thermo Fisher Scientific). The region flanking the gRNA targeting *ZEB1* was amplified using Taq DNA polymerase (Thermo Fisher Scientific) with primers as indicated in **Table S3**. PCR products were separated using a 1.5% agarose gel and DNA of specific bands was extracted using the Bioke Gel and PCR cleanup kit (Macherey-Nagel). Cloning of purified PCR products was performed using the InsTA PCR cloning kit (Thermo Fisher Scientific) and subsequently transformed in DH5α cells. For each PCR band identified in the agarose gel electrophoresis, 10 colonies were picked and grown overnight at 37 °C. Finally, plasmids were isolated using the Bioke plasmid cleanup kit (Macherey-Nagel). Samples were Sanger sequenced (Macrogen, Amsterdam, the Netherlands) using the M13 reverse primer (**Table S3**), after which sequencing results were analyzed using Snapgene version 7.2.1.

### Chromatin immunoprecipitation (ChIP) analysis

For chromatin immunoprecipitation (ChIP) assays, hPS1 ZEB1 KO cells, rescued with *ZEB1* cDNA (clone 2D12+ZEB1) were cultured for 48 hours. One day prior to fixation of the cells, Protein A dynabeads (Thermo Fisher Scientific) were coated with 4 µg ZEB1 (Proteintech 21544-1-AP, Manchester, UK) or 4 µg Rabbit IgG control Ab (PP64, Sigma-Aldrich) overnight in PBS/1%BSA (>98% BSA free, Sigma-Aldrich). Cells were fixed by adding 1% formaldehyde (Pharmacy LUMC) for 10 min, followed by quenching using 1/20 volume 2,5M glycine. After washing twice with PBS, cells were scraped from the plate and lysed using lysis buffer (50 mM Tris-HCl pH 8, 10 mM EDTA, 1% SDS and EDTA-free protease inhibitors (Sigma-Aldrich)). Samples were sonicated using the Bioruptor Pico (Diagenode, Seraing, Belgium), followed by centrifugation at 11,000 RPM for 10 min to clear debris.

5% input sample was saved, whereafter the remaining sample was diluted in dilution buffer (20 mM Tris-HCl pH 8, 2 mM EDTA, 1% Triton-X100, 150 mM NaCl and protease inhibitors). Antibody-bound beads were washed twice and added to the lysate, which was incubated O/N. The following day, samples were washed 5x in RIPA wash buffer (50 mM HEPES-KOH pH 7, 0.5M LiCl, 1 mM EDTA, 0.7% DOC and 1% Igepal) and once in TE buffer. 200 µl lysis buffer was added to the beads, followed by vortexing every 2 minutes for 15 minutes at 65°C and incubation at 65°C for 6 hours. DNA was purified using the Qiaquick PCR purification kit (Qiagen) and analyzed by qPCR using primers binding in the *F11R* or E-Cadherin promotor region (**Table S3**).

### Western blot

Total cell lysates were generated in Pierce RIPA buffer (Thermo Fisher Scientific), supplemented with complete mini protease inhibitor cocktail (Roche Applied Science, Penzberg, Germany). Samples were cleared from cellular debris by centrifugation (13,000 rpm, 4°C, 5 minutes). Protein concentrations were measured by the Pierce^TM^ BCA kit (Thermo Fisher Scientific). Lysates were denatured by adding Laemmli sample buffer containing 20 mM DTT and heating for 3 min at 95°C. Equal amounts of protein were separated by gel electrophoresis on 10% SDS-polyacrylamide gels and transferred onto 0.2 μm nitrocellulose membranes using the Trans-Blot Turbo Transfer System (Bio-Rad). Membranes were blocked in TBS, supplemented with 0.1% Tween20 (TBST) and 10% milk. Antibodies were diluted in TBST containing 5% milk. Primary antibodies were incubated overnight at 4°C, and secondary antibodies for 60 minutes at room temperature. Blots were washed with TBST.

The following primary antibodies were used: mouse anti-vinculin (Sigma-Aldrich, V9131), rabbit anti-ZEB1 (D80D3, Cell signaling, Leiden, the Netherlands) and mouse anti-reovirus σ3 (4F2, Developmental Studies Hybridoma Bank, developed under the auspices of the NICHD and maintained by the University of Iowa, Department of Biology, Iowa City, IA, USA).^(40)^ Proteins were visualized using the Odyssey CLx Imaging System (LI-COR Biosciences, Bad Homburg, Germany).

### Cell viability assays

To determine cell viability following reovirus infection, WST-1 reagent was employed (Roche, Woerden, the Netherlands). Cells were plated in 96 well plates (5-6 wells per condition) and infected with different MOIs of reovirus the following day. After 2-6 days, 20x diluted WST-1 reagent in infection medium was added to the wells. Absorption at OD450 was measured using a plate reader (ENZ-INS-A96, Enzo Life Sciences, Brussels, Belgium) and the percentages of cell viability were calculated by dividing the OD450 values of the virus-treated wells by the values of the mock condition.

### Caspase assays

To detect caspase 3/7 activation following reovirus infection, the CellEvent™ Caspase-3/7 green detection reagent (Thermo Fisher Scientific) was used. Cells were plated in triplicate in 96-well plates and infected with reovirus R124 at MOI 10 the following day. Phase contrast images, GFP images and quantifications of the GFP signal (excitation 500 nm, emission 530 nm) were taken every 30 min for a maximum of 48 hours using a Cytation Microplate Reader (Biotek). Images were analyzed using Fiji version 2.14.0.

### Statistical analysis

Data are presented as means ± standard deviation from representative experiments of independent replicates. Differences between more than 2 groups were measured using 1-way analysis of variance (ANOVA) or 2-way ANOVA, depending on the number of variables, and corrected for multiple testing. All analyses were performed using GraphPad Prism version 10.2.3 (San Diego, CA, USA). P values of 0.05 or less were considered statistically significant.

## Supporting information

Full data CRISPR screen replicate 1

Full data CRISPR screen replicate 2

## Data availability statement

Raw sequencing data can be accessed on Gene Expression Omnibus (GEO) under accession number GSE290358. The processed list of genes found from the CRISPR/Cas9 screen can be found in the supplemental data. All other data is included in the manuscript or supplemental data.

## Acknowledgements

Part of this work was carried out at the CRISPR Functional Genomics facility (CFG) at Karolinska Institutet funded by Science for Life Laboratory. CFG acknowledges support from the Swedish National Genomics Infrastructure, the National Academic Infrastructure for Supercomputing in Sweden (NAISS), and the Uppsala Multidisciplinary Center for Advanced Computational Science (UPPMAX). We further thank H. Kocher for providing us with the hPS1 cell line. The graphical abstract and Figure 1A were created using Biorender.com.

ND is sponsored by funding from the foundation “Overleven met Alvleesklierkanker” (Leiden, the Netherlands) (SOAK 21.02), obtained by VK. TJ Harryvan is sponsored by a personal MD-PhD grant from the Leiden University Medical Center and received personal funding from the Dutch Society for Medical Oncology to perform the genome-wide CRISPR/Cas9 KO screen at the Karolinska Institutet. VK is supported by a personal grant from the Dutch Research Council (NWO-talent program Veni, ZonMw).

## Author contributions

ND and TJH planned and performed the majority of the experiments and performed data analysis. TJH and BS performed the CRISPR/Cas9 KO screen. HD performed analysis of the CRISPR/Cas9 KO screen. GI performed DNA sequencing of the ZEB1 KO clones. VK and LJACH supervised the project. ND, TJH, LJACH and VK co-wrote the manuscript. All authors contributed to critically reviewing the manuscript.

