## Supplemental tables and figures +legends for "ZEB1, a novel regulator of Junctional Adhesion Molecule A, impacts sensitivity of pancreatic cancer-associated fibroblasts to oncolytic reovirus"

**Table S1. Sequences used during the genome-wide CRISPR/Cas9 KO screen**

| Name | Sequence |
|---|---|
| Oligo pool | CTTGTGGAAAGGACGAAACACCGNNNNNNNNNNNNNNNNNNNNGTTTAAGAGCT AGAAATAGCAAGTTTAAATAAGGCT |
| ds_Ultramer | TTTGTCTCAAGATCTAGTTACGCCAAGCTTNNNNNNNNNNGTGACTGGAGTTCAGA CGTGTGCTCTTCCGATCAAAAAAGCACCGACTCGGTGCCACTTTTTCAAGTTGATAAC GGACTAGCCTTATTTAAACTTGCTATTTCTAGCTC |
| ds_fw | GGCTTTATATATCTTGTGGAAAGGACGAAACACCG |
| ds_rev | TTTGTCTCAAGATCTAGTTACGCCAAGC |
| Final insert | GGCTTTATATATCTTGTGGAAAGGACGAAACACCGNNNNNNNNNNNNNNNNNNN NGTTTAAGAGCTAGAAATAGCAAGTTTAAATAAGGCTAGTCCGTTATCAACTTGAAA AAGTGGCACCGAGTCGGTGCTTTTTTGATCGGAAGAGCACACGTCTGAACTCCAGTC ACNNNNNNNNNNAAGCTTGGCGTAACTAGATCTTGAGACAAA |
| PCR1_fw | GGACTATCATATGCTTACCGTAACTTGAAAGTATTTCG |
| PCR1_rev | CTTTAGTTTGTATGTCTGTTGCTATTATGTCTACTATTCTTTCC |
| PCR2_fw | ACACTCTTTCCCTACACGACGCTCTTCCGATCTCTTGTGGAAAGGACGAAACAC |
| PCR2_rev | AGAAGACGGCATACGAGATCTGCCATTTGTCTCAAGATCTAGTTAC |
| PCR3_fw | AATGATACGGCGACCACCGAGATCTACAC[I5]ACACTCTTTCCCTACACGACGCTCT |
| PCR3_rev | CAAGCAGAAGACGGCATACGAGATCTGCCATTTG |
| CRISPRSeq | CGATCTCTTGTGGAAAGGACGAAACACCG |

**Table S2. Mission shRNA constructs**

| Mission shRNA constructs | Sequence (5'-3') |
|---|---|
| Non-targeting control SHC002 | CCGGCAACAAGATGAAGAGCACCAACTCG-<br>AGTTGGTGCTCTTCATCTTGTTGTTTTT |
| Mouse ZEB1 KD#1 (TRCN0000235850) | ATAGAGGCTACAAGCGCTTTA |
| Mouse ZEB1 KD#2 (TRCN0000235853) | GTCGACAGTCAGTAGCGTTTA |
| Mouse FGFR1 KD#1 (TRCN0000023295) | CCTGGAGCATCATAATGGATT |
| Human ZEB1 KD#1 (TRCN0000017565) | CCTCTCTGAAAGAACACATTA |
| Human ZEB1 KD#2 (TRCN0000364631) | CCTACCACTGGATGTAGTAAA |
| Human FGFR1 KD#1 (TRCN0000312574) | TGCCACCTGGAGCATCATAAT |
| Human FGFR1 KD#2 (TRCN0000121185) | CCACAGAATTGGAGGCTACAA |
| Human SPPL3 KD#1 (TRCN0000307051) | CCTGGTCTCCTACTATGCTTT |
| Human SPPL3 KD#2 (TRCN0000308107) | GGGCATCGGAGACATCGTTAT |

**Table S3. Primer sequences**

| Gene | Primer sequence (5’-3’) |
|---|---|
| M13 Reverse sequencing primer | CAGGAAACAGCTATGAC |
| *ZEB1* KO validation | Fw: CACCACACCTGAGGAAAAC<br>Rv: TTTCCCACTCCACTTTGCCGTC |
| ChIP *JAM-A* 1 | Fw: GCCTGCAACATCTCCCGTT<br>Rv: ATGTTAAGGGCTTCTGCGGTG |
| ChIP *JAM-A* 2 | Fw: ACAGGAGCTGCCTCAGATTGG<br>Rv: GTACTTCTCAGCCCTCTAGCTC |
| ChIP *JAM-A* 3 | Fw: ACTACAGCGAGGGGACTGAG<br>Rv: GAAGAGCAGCGGTTCCTTAC |
| ChIP *JAM-A* 4 | Fw: TGCACGTTCCGATTGGTGTAA<br>Rv: AGTCTCCTGGGCCAATCTGAG |
| ChIP *E-Cadherin* | Fw: GGCCGGCAGGTGAAC<br>Rv: GGGCTGGAGTCTGAACTGAC |
| *ACTB* (mouse) | Fw: AGGTCATCACTATTGGCAACGA<br>Rv: CCAAGAAGGAAGGCTGGAAAA |
| *Mzt2* (mouse) | Fw: TCGGTGCCCATATCTCTGTC<br>Rv: CTGCTTCGGGAGTTGCTTTT |
| *Ptp4a2* (mouse) | Fw: AGCCCCTGTGGAGATCTCTT<br>Rv: AGCATCACAAACTCGAACCA |
| *ZEB1* (mouse) | Fw: ATTCAGCTACTGTGAGCCCTGC<br>Rv: CATTCTGGTCCTCCACAGTGGA |
| *FGFR1* (mouse) | Fw: GCCTCACATTCAGTGGCTGAAG<br>Rv: AGCACCTCCATTTCCTTGTCGG |
| *JAM-A* (mouse) | Fw: CACCTACTCTGGCTTCTCCTCT<br>Rv: TGCCACTGGATGAGAAGGTGAC |
| *IPO8* (human) | Fw: AGGATCAGAGGACAGCACTGCA<br>Rv: AGGTGAAGCCTCCCTGTTGTTC |
| *EIF2B1* (human) | Fw: CTACTCCAGAGTGGTCCTGAGA |

| | |
|---|---|
| | Rv: GTTGAGGTGGCAGAGGGCTTTG |
| *JAM-A* (human) | Fw: GTGAAGTTGTCCTGTGCCTACTC<br>Rv: ACCAGTTGGCAAGAAGGTCACC |
| *ZEB1* (human) | Fw: GGCATACACCTACTCAACTACGG<br>Rv: TGGGCGGTGTAGAATCAGAGTC |
| *FGFR1* (human) | Fw: GCACATCCAGTGGCTAAAGCAC<br>Rv: AGCACCTCCATCTCTTTGTCGG |

### Supplemental figures

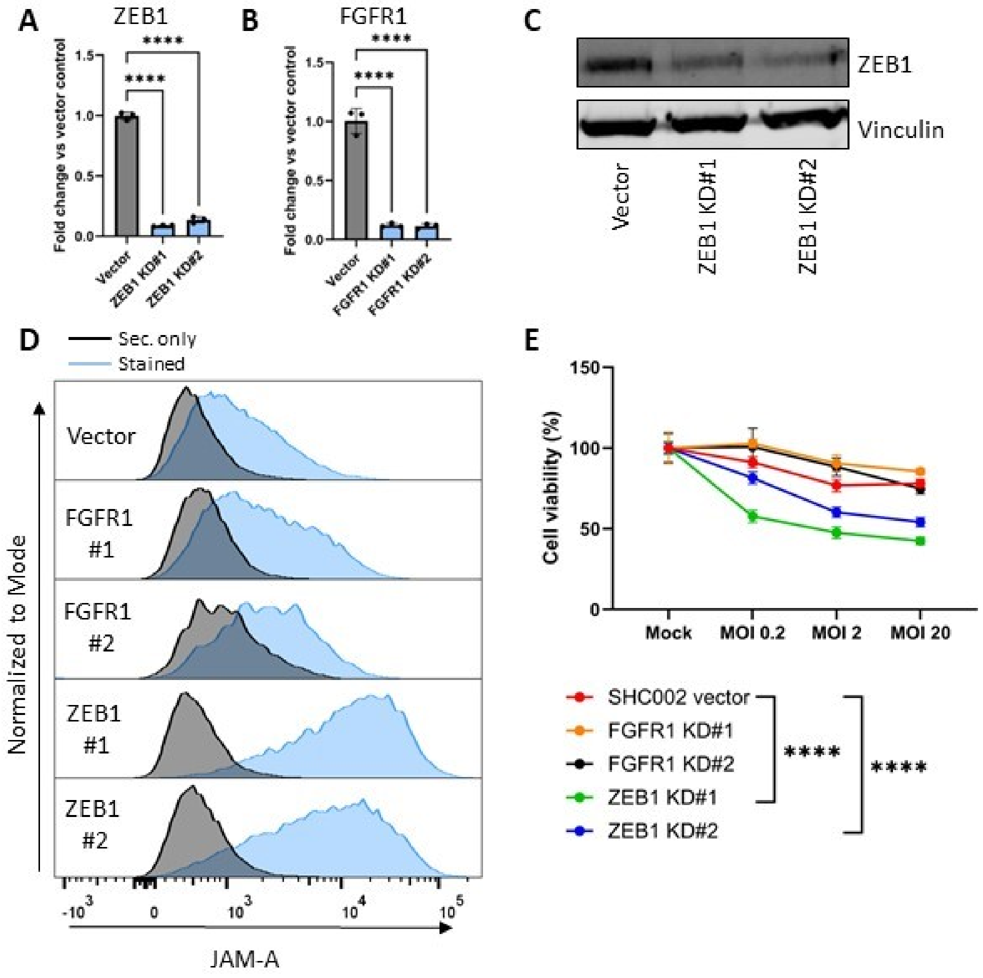


**Figure S1.** Knockdown of ZEB1, but not FGFR1, in a skin fibroblast cell line NBS results in JAM-A upregulation and sensitization to reovirus-mediated cell death. **A,B** RT-qPCR for *ZEB1* in NBS vector control and ZEB1 KD (A) and *FGFR1* in NBS vector control and FGFR1 KD (B). Ct values were corrected for *β-actin* expression and calculated as fold change vs vector control. ****p≤0,0001 as determined by one-way ANOVA with correction for multiple testing (Šídák's test). Data is derived from a representative experiment and plotted as mean ± SD. **C** Western blot for ZEB1 with vinculin as loading control. **D** Flow cytometric analysis of cell-surface JAM-A expression in NBS vector control and FGFR1 and ZEB1 KD. Black: secondary antibody only, blue: stained. **D** Cell viability (%) relative to mock following infection with reovirus at multiple MOIs for 5 days, as measured by a WST-1 assay. Significance was calculated using two-way ANOVA with correction for multiple testing (Šídák's test), significance is depicted at MOI 10, ****p≤0,0001. Data is derived from a representative experiment and plotted as mean ± SD.

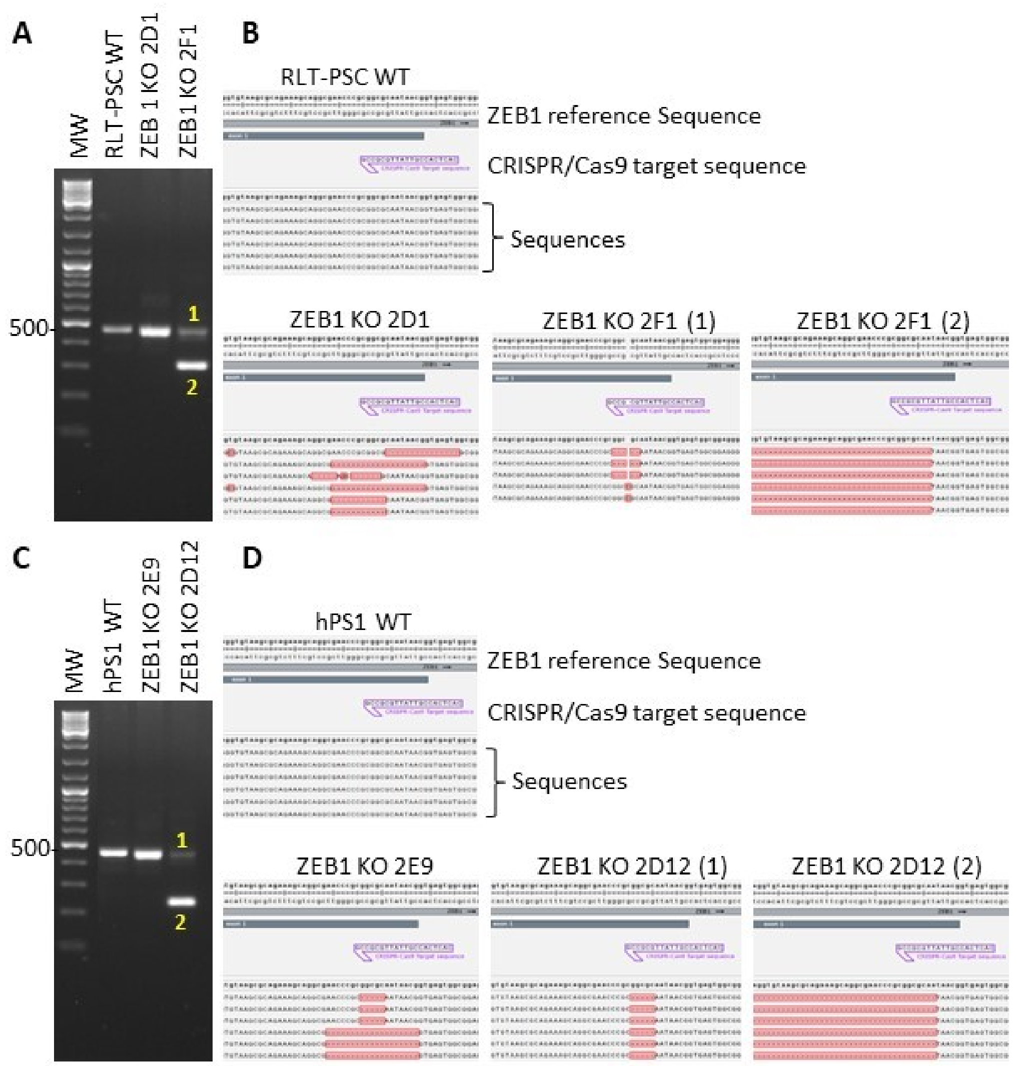


**Figure S2.** Sequencing of *ZEB1* in the region targeted by the gRNA shows KO-responsible indels. **A** Agarose gel electrophoresis following PCR of RLT-PSC WT and ZEB1 KO clones 2D1 and 2F1. **B** Sanger sequence results of RLT-PSC WT and ZEB1 KO 2D1 and 2F1 (2 different bands) following isolation from the agarose gel, ligation into a vector and sanger sequencing. The different sequences shown are the result of the different colonies isolated from the miniprep. **C** Agarose gel electrophoresis following PCR of hPS1 WT and ZEB1 KO clones 2E9 and 2D12. **D** Sanger sequence results of hPS1 WT and ZEB1 KO 2E9 and 2D12 (2 different bands) following isolation from the agarose gel, ligation into a vector and sanger sequencing. The different sequences shown are the result of the different colonies isolated from the miniprep.

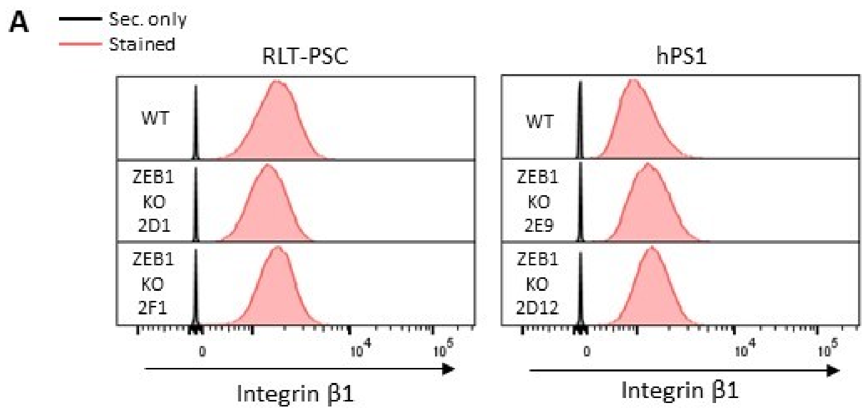


**Figure S3.** Integrin β1 expression is unchanged following ZEB1 KO. **A** Flow cytometric analyses of Integrin β1 expression in RLT-PSC WT and ZEB1 KO (left panel) and hPS1 WT and ZEB1 KO (right panel). Black: secondary antibody only, red: stained.

### Supplemental video legend

**Supplemental video 1.** CellEvent Caspase 3/7 assay in RLT-PSC, hPS1 and KPC3-CAF1 WT and ZEB1 KO following infection with reovirus R124 MOI 10 for 36, 48 and 24 hours, respectively. Phase-contrast and GFP images are overlayed and made into videos using Fiji version 2.14.0. Scale bar: 200 µM.
